# Genome-scale reconstructions of the mammalian secretory pathway predict metabolic costs and limitations of protein secretion

**DOI:** 10.1101/351387

**Authors:** Jahir M. Gutierrez, Amir Feizi, Shangzhong Li, Thomas B. Kallehauge, Hooman Hefzi, Lise M. Grav, Daniel Ley, Deniz Baycin Hizal, Michael J. Betenbaugh, Bjorn Voldborg, Helene Faustrup Kildegaard, Gyun Min Lee, Bernhard O. Palsson, Jens Nielsen, Nathan E. Lewis

**Affiliations:** Department of Bioengineering, University of California, San Diego, La Jolla, CA 92093, United States; Novo Nordisk Foundation Center for Biosustainability at the University of California, San Diego, School of Medicine, La Jolla, CA 92093, United States; Department of Biology and Biological Engineering, Kemivägen 10, Chalmers University of Technology, SE41296 Gothenburg, Sweden; Novo Nordisk Foundation Center for Biosustainability, Technical University of Denmark, 2800 Kgs. Lyngby, Denmark; Department of Systems Biology, Technical University of Denmark, Kongens Lyngby, Denmark; Pharmaceutical R&D Department, Turgut Illaclari A.S., Istanbul, Turkey; Department of Chemical & Biomolecular Engineering, Johns Hopkins University, Baltimore, MD 21218-2686, United States; Department of Pediatrics, University of California, San Diego, School of Medicine, La Jolla, CA 92093, United States

**Keywords:** Metabolic network, secretory pathway, biotherapeutic production, systems biotechnology

## Abstract

In mammalian cells, >25% of synthesized proteins are exported through the secretory pathway. The pathway complexity, however, obfuscates its impact on the secretion of different proteins. Unraveling its impact on diverse proteins is particularly important for biopharmaceutical production. Here we delineate the core secretory pathway functions and integrate them with genome-scale metabolic reconstructions of human, mouse, and Chinese hamster cells. The resulting reconstructions enable the computation of energetic costs and machinery demands of each secreted protein. By integrating additional omics data, we find that highly secretory cells have adapted to reduce expression and secretion of other expensive host cell proteins. Furthermore, we predict metabolic costs and maximum productivities of biotherapeutic proteins and identify protein features that most significantly impact protein secretion. Finally, the model successfully predicts the increase in secretion of a monoclonal antibody after silencing a highly expressed selection marker. This work represents a knowledgebase of the mammalian secretory pathway that serves as a novel tool for systems biotechnology.

## Introduction

To interact with their environment, cells produce numerous signaling proteins, hormones, receptors, and structural proteins. In mammals, these include at least 2,641 secreted proteins (e.g., enzymes, hormones, antibodies, extracellular matrix proteins) and >5,500 membrane proteins^1^, most of which are synthesized and processed in the secretory pathway.

The secretory pathway consists of a complex series of processes that predominantly take place in the endoplasmic reticulum (ER), Golgi apparatus, and the endomembrane system. This pathway is particularly important in biotechnology and the biopharmaceutical industry, since most therapeutic proteins are produced in mammalian cell lines such as HEK293, PerC6, NS0, and Chinese hamster ovary (CHO) cells, which are capable of folding and adding the necessary post-translational modifications (PTMs) to the target product^2^. For any given biotherapeutic, different machinery in the secretory pathway may be needed, and each step can exert a non-negligible metabolic demand on the cells. The complexity of this pathway, however, makes it unclear how the biosynthetic cost and cellular needs vary for different secreted proteins, each of which exerts different demands for cellular resources. Therefore, a detailed understanding of the biosynthetic costs of the secretory pathway could guide efforts to engineer host cells and bioprocesses for any desired product. The energetic and material demands of the mammalian secretory pathway can be accounted for by substantially extending the scope of metabolic models. Indeed, recent studies have incorporated portions of the secretory pathway in metabolic models of yeast ^3–5^. Furthermore, Lund and colleagues reconstructed a genetic interaction network of the mouse secretory pathway and the unfolded protein response and analyzed it in the context of CHO cells^6^. However, such a network does not encompass a stoichiometric reconstruction of the biochemical reactions involved in the secretory pathway nor it is coupled to existing metabolic networks of mammalian cells.

Here we present the first genome-scale stoichiometric reconstructions and computational models of mammalian metabolism coupled to protein secretion. Specifically, we constructed these for human, mouse, and CHO cells, called RECON2.2s, iMM1685s, and iCHO2048s, respectively. We first derive an expression for computing the energetic cost of synthesizing and secreting a product in terms of molecules of ATP equivalents per protein molecule. We use this expression and analyze how the energetic burden of protein secretion has led to an overall suppression of more expensive secreted host cell proteins in mammalian cells. Given its dominant role in biotherapeutic production, we further focus on the biosynthetic capabilities of CHO cells. We then demonstrate that product-specific secretory pathway models can be built to estimate CHO cell growth rates given the specific productivity of the recombinant product as a constraint. We identify the features of secreted proteins that have the highest impact on protein cost and productivity rates. Finally, we use our model to identify proteins that compete for cell resources, thereby presenting targets for cell engineering. Through this study we demonstrate that a systems-view of the secretory pathway now enables the analysis of many biomolecular mechanisms controlling the efficacy and cost of protein expression in mammalian cells. We envision our models as valuable tools for the study of normal physiological processes and engineering cell bioprocesses in biotechnology. All models and data used in this study are freely available at https://github.com/LewisLabUCSD/MammalianSecretoryRecon.

## RESULTS

### A stoichiometric expression of protein secretion energetics

In any cell, the secretory machinery is concurrently processing thousands of secreted and membrane proteins, which all compete for secretory pathway resources and pose a metabolic burden. To quantify this burden, we estimated the energetic cost of synthesizing and/or secreting 5,641 and 3,538 endogenous proteins in the CHO and human secretome and membrane proteome in terms of total number of ATP equivalent molecules consumed (see Methods). These protein costs were compared to the cost of five recombinant proteins commonly produced in CHO cells (Fig. 1a). To refine estimates, we predicted signal peptides^7^, GPI anchor attachment signals^8^, and experimentally measured the number of N-linked glycans in the CHO proteome and integrated published numbers of O-linked glycans in CHO proteomic data^9^. Across the CHO secretome, protein synthesis cost varies substantially, and recombinant products are on average more expensive (Fig. 1a). For example, Factor 8 (F8) is a difficult-to-express protein in CHO cells due to its propensity to aggregate in the ER, which promotes its premature degradation^10,11^. Our analysis further highlights that each molecule of F8 requires a large amount of ATP for its production (9,488 ATP molecules). This imposes a significant burden to the secretory machinery of CHO cells, which typically expresses much less expensive endogenous proteins.

**Figure 1.**
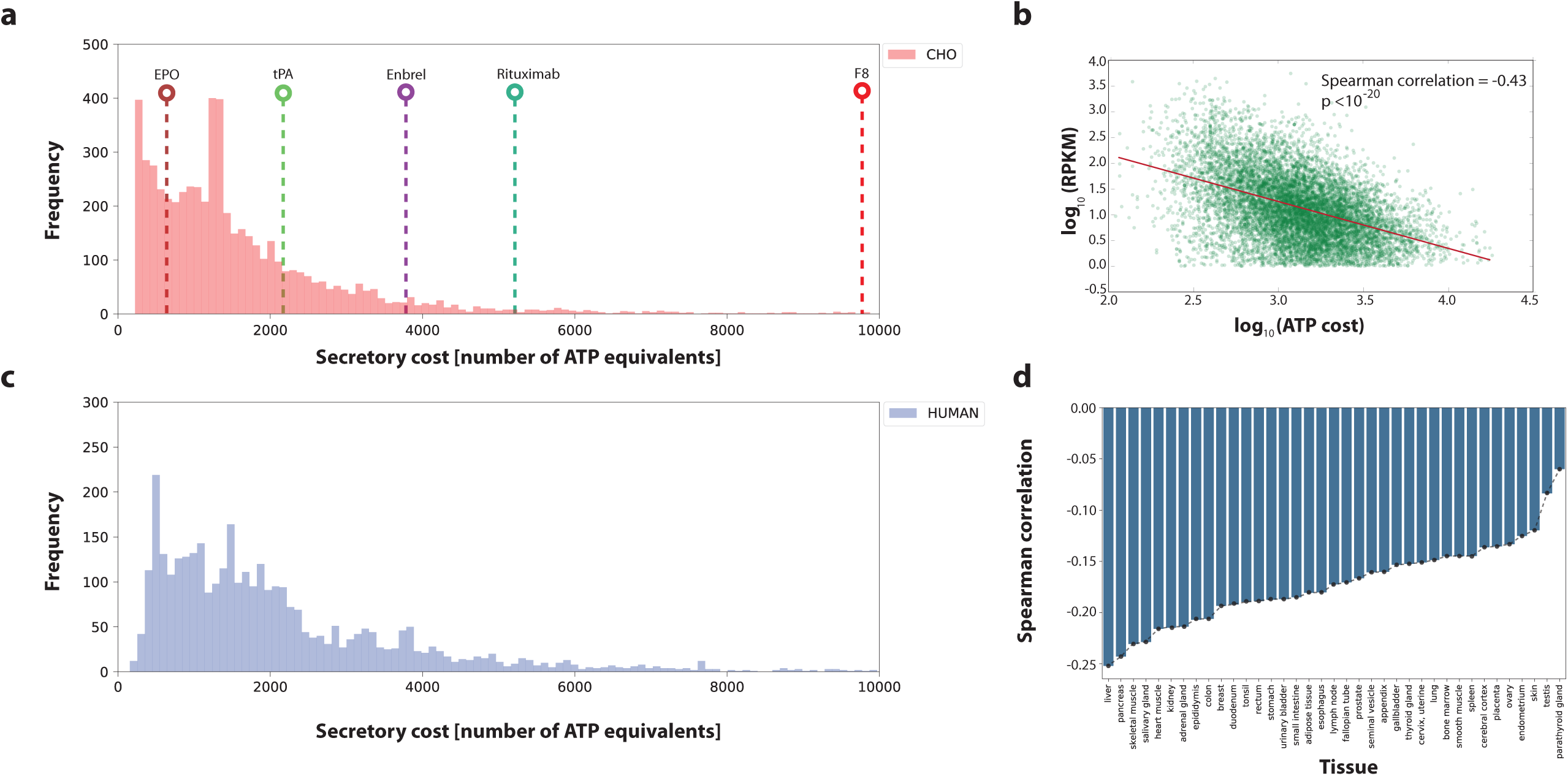
Mammalian secretory cells preferentially suppress more expensive proteins. The bioenergetic cost of each secreted CHO (a) and human (c) protein was computed. The bioenergetic costs of five representative biotherapeutics produced in CHO cells are shown for comparison purposes (see Table 1). (b) Scatter plot and Spearman correlation of gene expression measured by ribosomal profiling and protein cost (in number of ATP per protein) in CHO cells from Kallehauge et al.^12^ during the early exponential growth phase of culture. (d) Spearman correlations between ATP cost and gene expression levels (measured by RNA-seq) across human tissues^1, 58^. Gene transcription levels from the Human Protein Atlas were analyzed against the ATP cost of producing the translated proteins. All p-values associated to each correlation are <1×10^-20^. Highly secretory tissues show the strongest negative correlation of secreted protein cost vs. mRNA expression levels. RPKM = reads per kilobase of transcript per million. Source data are provided as a Source Data file.

### Recombinant cells suppress expression of expensive proteins

With the broad range of biosynthetic costs for different proteins, we wondered if gene expression in mammalian cells that are tasked with high levels of protein secretion have been influenced by the ATP cost of secreted proteins. That is, have these secretory cells suppressed their protein expression to more efficiently allocate nutrients? To test this, we first looked at CHO cells, which have undergone extensive selection to obtain cells that secrete recombinant proteins at high titer, and then compared different human tissues with a range of secretory capacity.

**Table 1.** Protein specific information matrix of biotherapeutics secreted in eight iCHO2048s models.

| Protein Name | Length [AA] | Weight [Da] | Disulfide bonds | N-glycans | O-glycans | ATP cost |
| --- | --- | --- | --- | --- | --- | --- |
| IFNB1 | 187 | 22294 | 1 | 1 | 0 | 777 |
| EPO | 193 | 21037 | 2 | 3 | 1 | 801 |
| BMP2 | 396 | 44702 | 4 | 5 | 0 | 1618 |
| BMP7 | 431 | 49313 | 4 | 4 | 0 | 1759 |
| tPA | 562 | 61917 | 17 | 3 | 1 | 2286 |
| Etanercept* | 934 | 102470 | 7 | 6 | 26 | 3784 |
| Rituximab** | 1328 | 143860 | 17 | 2 | 0 | 5370 |
| F8 | 2351 | 267009 | 8 | 22 | 0 | 9488 |

\* = *Etanercept is a dimer*
\*\*= *Rituximab is a tetramer (2 light and 2 heavy chains)*

Unless specific proteins are essential, CHO cells may preferentially suppress energetically expensive proteins. Thus, we analyzed ribosomal profiling (Ribo-Seq) data from a recombinant CHO cell line^12^ and compared translation of each transcript against the ATP cost of the associated secreted protein (see Methods). Indeed, there was a significant negative correlation of −0.43 (Spearman R_s_, p value < 1×10^-20^) between ribosomal occupancy and ATP cost during early exponential growth phase of culture (Fig. 1b). Wondering if the reduced translation was regulated transcriptionally, we further analyzed RNA-Seq data from the same recombinant cell line and from another, non-recombinant CHO-K1 cell line^13^. The RNA expression also negatively correlated with ATP cost (see Supplementary Figure 2).

To evaluate if this is a general trend seen in mammalian secretory cells, we analyzed RNA-Seq data from human tissues and immortalized cell lines in the Human Protein Atlas (HPA)^1^. For all RNA-Seq datasets in the HPA, there was a negative correlation between mRNA expression levels and ATP cost (Fig. 1D). Interestingly, we found that highly secretory tissues such as liver, pancreas and salivary gland had the strongest correlations, although none as strong as that of the recombinant CHO cells, which have undergone selection of high secretion. Feizi and colleagues recently found that these tissues fine-tune the expression of protein disulfide isomerase genes^14^, suggesting that a similar regulatory process may take place in the ER of CHO cells as the secreted monoclonal antibody (mAb) contains a relatively high number (17) of disulfide bonds. In conclusion, there is a clear preference in CHO and native secretory tissues to suppress the expression and translation of proteins that are costly to synthesize, fold, and secrete.

### In silico reconstruction of the mammalian secretory pathway

We mapped out the core processes involved in the synthesis of secreted and membrane proteins in mammalian cells (i.e., human, mouse, and Chinese hamster). This included 261 components (gene products) in CHO cells and 271 components in both human and mouse. The components are involved in secretory reactions across 12 subsystems (i.e., functional modules of the secretory pathway; Fig. 2a). These components represent the core secretory machinery needed in the transition of a target protein from its immature state in the cytosol (i.e., right after translation) to its final form (i.e., when it contains all post-translational modifications and is secreted to the extracellular space). Each component in the reconstruction either catalyzes a chemical modification on the target protein (e.g., N-linked glycosylation inside ER lumen/Golgi) or participates in a multi-protein complex that promotes protein folding and/or transport. This distinction between catalytic enzymes and complex-forming components is important for modeling purposes as a catalytic component consumes or produces metabolites that are directly connected to the metabolic network (e.g., ATP, sugar nucleotides). Because all components of the core secretory pathway were conserved across human, mouse and hamster (Fig. 2b), we generated species-specific secretory pathway reconstructions and used them to expand the respective genome-scale metabolic networks (Recon 2.2^15^, iMM1415^16^, iCHO1766^17^). Following the naming convention of their metabolic counterparts, we named these new metabolic-secretory reconstructions as follows: iMM1685s, iCHO2048s, and Recon 2.2s, which account for 1685, 2048, and 1946 genes, respectively. A detailed list of the components, reactions and the associated genes can be found in the Supplementary Data 1.

**Figure 2.**
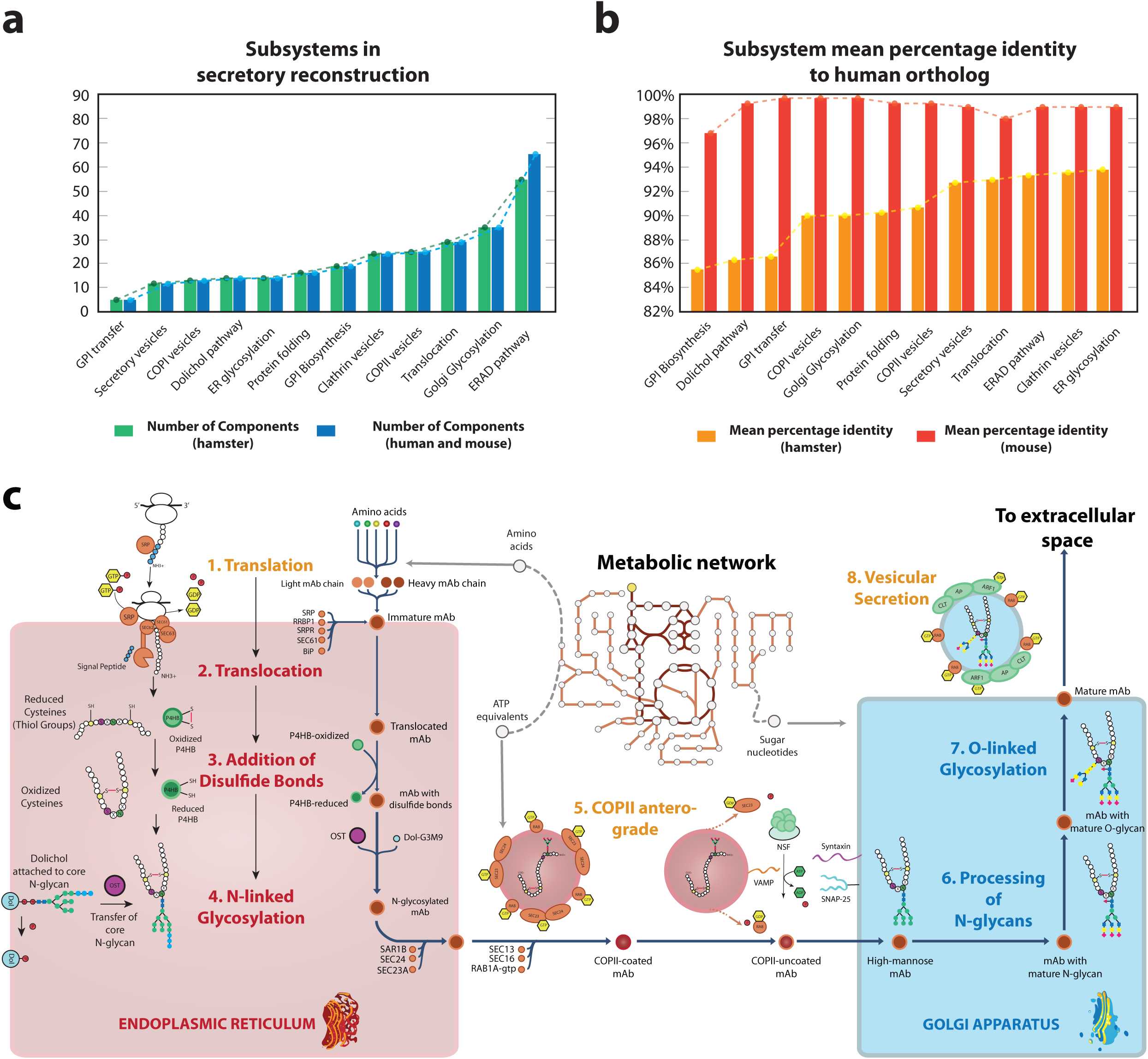
Components in the reconstruction of the secretory pathway in mammalian cells. (a) The reconstruction comprises 261 proteins in CHO cells and 271 proteins in human and mouse that are distributed across 12 subsystems. The different component numbers arise from the fact that the Chinese hamster proteome annotation only contains one alpha and one beta proteasome subunits, whereas the human and mouse contain 12 subunits of different subtypes. (b) High similarities were seen for proteins in CHO and human, with a high mean percentage identity in each subsystem (calculated with the sequence alignment tool BLAST). (c) Simplified schematic of reactions and subsystems involved in the secretion of a monoclonal antibody (mAb). A total of eight subsystems are necessary to translate, fold, transport, glycosylate, and secrete a mAb. The color of the subsystem names indicates if the reactions occur in the cytoplasm (orange), the ER lumen (red) or the Golgi apparatus (blue). The detailed description of all components can be found in Supplementary Data 1. GPI = Glycosylphosphatidylinositol, ER = Endoplasmic Reticulum, ERAD = ER associated degradation.

### Validation of iCHO2048s growth and productivity predictions

We first validated the accuracy of iCHO2048s predictions using growth and specific productivity rates of IgG-producing CHO cell lines from two independent studies^12,18^. For this, we built an IgG-secreting iCHO2048s model using the information in the PSIM matrix for the therapeutic mAb Rituximab. We then constrained the model’s Rituximab-specific secretory pathway with the reported productivity value in each study and used FBA to predict growth (Fig. 3a). Later, to assess the ability of iCHO2048s to predict growth rates in cases when CHO cells are producing non-antibody proteins, we collected data from two batch culture experiments using Enbrel- and C1-inhibitor-producing isogenic CHO cell lines. We constructed two iCHO2048s models for each case and predicted growth rates during the early exponential growth phase of culture while constraining the protein secretion rate to the measured specific productivity value (Fig. 3b). The model predictions agreed well with the reported and measured values. There were cases where iCHO2048s predicted a much higher growth rate than what was measured in the first days of batch culture (Fig. 3b). Since FBA computes theoretical maximum growth rates given a set of constraints, these over-prediction cases point at situations where CHO cells do not direct resources towards biomass production (during very early stages of culture), a discrepancy that is attenuated in later stages of culture. In conclusion, these results confirm the ability of protein-specific reconstructions to capture the specific energetic requirements that each recombinant product imposes on CHO cell metabolism.

**Figure 3.**
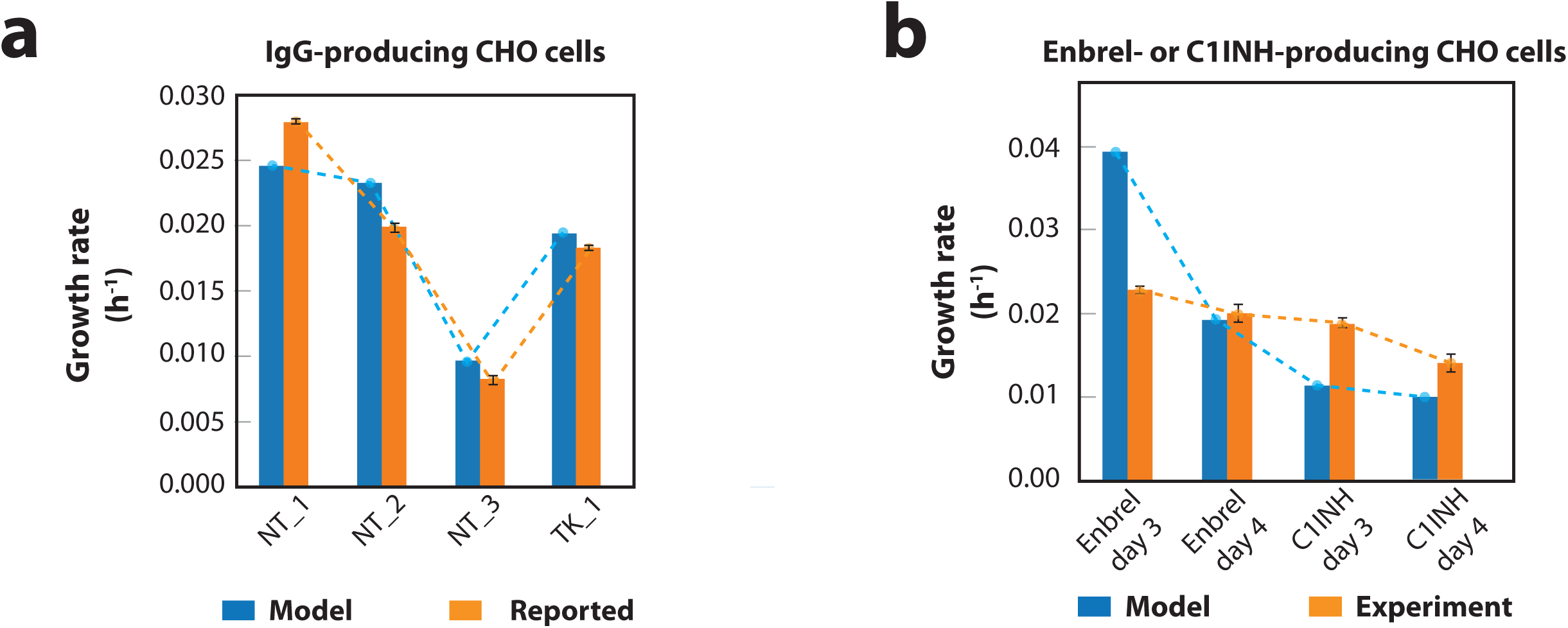
Recombinant-protein-producing models of iCHO2048s predict measured growth rates. (a) Growth rates were computed using an IgG-specific iCHO2048s model and compared to experimentally measured growth rates from six datasets from two previous studies using IgG-producing cell lines^12,18^. NT and TK specify the initials of the first author of the two studies (Neil Templeton, Thomas Kallehauge). (b) Additional growth, productivity, and metabolomic data were obtained from Enbrel and C1INH-producing CHO cells, and models were constructed. The model-predicted growth rates during exponential growth phase were consistent with experimental growth rates of Enbrel-producing CHO cells and C1INH-producing CHO cells at almost all time points. In all cases, the iCHO2048s models were constrained to produce the recombinant protein at the measured specific productivity rate. The values used to constrain each of the iCHO2048s models are reported in Supplementary Data 3. Error bars represent the standard deviation of three biological replicates. Source data are provided as a Source Data file.

### Protein composition impacts predicted productivity

To produce a specific product, CHO cells may utilize different modules of the secretory pathway based on the protein attributes and post-translational modifications (PTMs). For example, the synthesis of a mAb requires the use of multiple processes and consumes several different metabolites, such as amino acids for protein translation, redox equivalents for forming disulfide bonds, ATP equivalents for vesicular transport, and sugar nucleotides for protein glycosylation (Fig. 2c). Therefore, we generated eight product-specific secretory pathway models for biotherapeutics commonly produced in CHO cells (Fig. 4a): bone morphogenetic proteins 2 and 7 (BMP2, BMP7), erythropoietin (EPO), Enbrel, factor VIII (F8), interferon beta 1a (IFNB1), Rituximab, and tissue plasminogen activator (tPA). The resulting iCHO2048s models were used to compute Pareto optimality frontiers between maximum cell growth (*μ*) and specific productivity (*q*_P_), given the same measured glucose and amino acid uptake rates for each model^17^ (see Supplementary Data 3).

**Figure 4.**
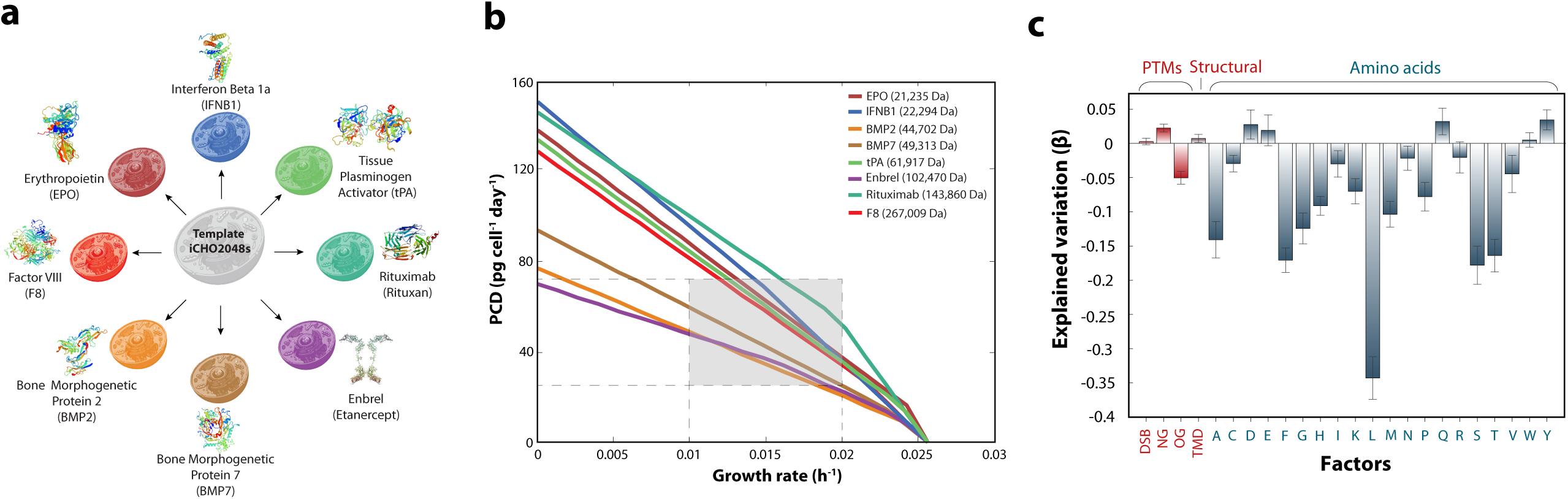
Construction of product-specific iCHO2048s models. (a) Eight product-specific iCHO2048s models were constructed for biotherapeutics commonly produced in CHO cells. (b) Pareto optimality frontiers of growth-productivity trade-off curves were computed for the eight iCHO2048s models using the same constraints and experimental data from Supplementary Data 3. The shaded region corresponds to range of maximum productivity at commonly observed growth rates in CHO cell cultures. The molecular weight (in Daltons) of each biotherapeutic is shown in the legend. (c) All protein features (PTMs, transmembrane domains, and amino acid compositions) were used to fit a multivariate linear regression to predict specific productivity. The model coefficients (β) quantify their contribution to the explained variation in specific productivity. Error bars represent the standard error of the fitted coefficients. Source data are provided as a Source Data file.

We computed the tradeoff between growth rate (hours^-1^) and specific productivity (picogram of protein produced per cell per day, or PCD) as a Pareto optimal curve for each protein (Fig. 4b). This curve defines the frontier of maximum specific productivity and maximum growth rates under the assumption that CHO cells can utilize all available resources towards production of biomass and recombinant protein only. The hinges in some of the curves are indicative of a transition between regions that are limited by distinct protein requirements (e.g., amino acids).

An analysis of the Pareto optimal curves for the eight biotherapeutics demonstrates that under the measured growth conditions, maximum productivities vary from 20-100 PCD at common growth rates (Fig. 4b, shaded region) to 70-150 PCD for senescent CHO cells. Neither the molecular weight (MW) nor product length can explain the 2-fold range differences in maximum productivity for different proteins. For example, the curves show tPA (MW = 61,917 Da) can express at higher PCD than BMP2 (MW = 44,702 Da) despite being larger, because the N-glycans in BMP2 reduce productivity due to the higher cost of synthesizing core N-glycans (see Table 1), consistent with previous observations in yeast^5^. Furthermore, the degree and directionality of these effects will depend on the nutrient uptake rates (Figure 4c and Supplementary Figure 1), highlighting the need in CHO bioprocessing to tailor culture media in a host cell and product-specific manner. Thus, while intuitively larger proteins would be expected to exert more bioenergetic cost on protein secretion, we find that specific compositional attributes of both the recombinant protein and the culture media significantly impact biosynthetic capacity. An in-depth analysis of the effects of PTMs on predicted productivities is provided in Jupyter Notebook C.

To further evaluate what functions of the secretory pathway had the greatest impact on the cost of protein synthesis and secretion, we computed secretion rates for 5,461 proteins in the CHO secretome (see Methods) using iCHO2048s and its parent metabolic reconstruction iCHO1766^17^. While iCHO2048s captures all the required steps for protein synthesis, modification and secretion, the secretion reactions in iCHO1766 only account for the basic synthesis of the target protein in cytoplasm, and the synthesis of necessary precursors (N-linked glycans, O-linked glycans, and GPI anchors). We found that secretory pathway had non-negligible costs on most proteins (Supplementary Figure 3b). Furthermore, protein features associated with secreted proteins that differ in cost by >15% beyond the amino acid and glycan costs show a statistical enrichment (under the Hypergeometric test) for O-linked glycans (p = 0.0065), GPI anchors (p = 0.0216), transmembrane domains (p = 0.0326), and proteins destined to the ER lumen (p=0.0142), the Golgi membrane (p = 0.0065), or the plasma membrane (p = 0.0186, see Supplementary Figure 3d and Jupyter Notebook E). Thus, these PTMs and transmembrane domains exert additional costs to their demands.

### iCHO2048s recapitulates results following gene knock-down

In a recent study, Kallehauge et al.^12^ demonstrated that a CHO-DG44 cell line producing an antiviral mAb^19^ also expressed high levels of the *neoR* selection-marker gene (Fig. 5a-b). Upon *neoR* knockdown, the titer and maximum viable cell densities of the CHO-DG44 cell line were increased. To test if iCHO2048s could replicate these results, we constructed a model for the Kallehauge et al. DG44 cell line and measured exometabolomics, and dry cell weight to parameterize the model. Since expression of *neoR* uses resources that could be used for antibody production, we predicted how much additional antibody could be synthesized with the elimination of the *neoR* gene. We simulated antibody production following a complete knockout of *neoR* (see Table 2 and Fig. 5b) and predicted that the deletion of *neoR* could increase specific productivity by up to 4% and 29% on days 3 (early exponential phase) and 6 (late phase) of culture, respectively (Fig. 5c). This was qualitatively consistent with the experimentally observed values of 2% and 14% when *neoR* mRNA was knocked down by 80-85%. We then computed the Pareto optimality curves for both the control and the *neoR* in silico knockout conditions on day 6. We found that the length of the curve (denoted by *Δ*) increased by 18% when *neoR* production is eliminated (Fig. 5d). Thus, iCHO2048s can quantify how much non-essential gene knockouts can boost growth and productivity in CHO cells by freeing energetic and secretory resources. In fact, the ribosome-profiling data from Kallehauge et al. revealed that only 30 secretory proteins in CHO cells account for more than 50% of the ribosomal load directed towards translation of protein bearing a signal peptide (Fig. 4E). Indeed, we recently found that substantial resources can be liberated and recombinant protein titers can be increased when 14 high-abundance host cell proteins were knocked out^20^. An analysis of other potential host cell gene knockouts using the method proposed here can be found in Supplementary Data 4.

**Table 2.** Experimental data used for validation of iCHO2048s predictive capabilities.

| Measurement [Units] | Early growth phase | Late growth phase |
| --- | --- | --- |
| Growth Rate [1 per day] | 0.44 | 0.02 |
| Specific Productivity [Picograms of IgG per cell perday]* | 16 | 5.5 |
| Total IgG ribosomal footprint [RPKM]** | 40258 | 13356 |
| Total neoR ribosomal footprint [RPKM] | 36952 | 25679 |

\* Average cell dry weight = 456.3 pg per cell
\*\* Sum of light and heavy chains ribosomal footprints

**Figure 5.**
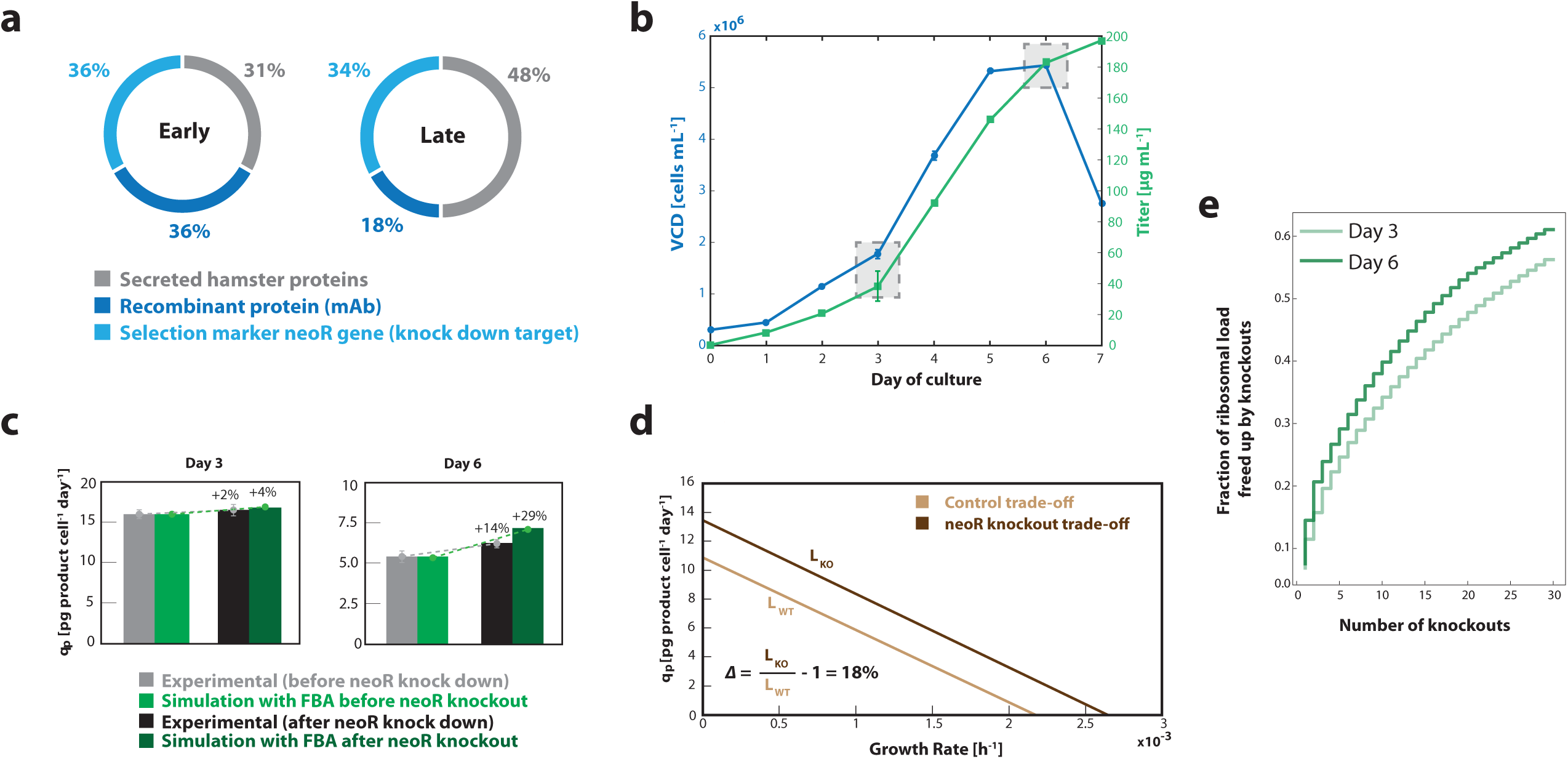
iCHO2048s recapitulates experimental results of *neoR* knock-down in silico. (a) Ribosome occupancy was measured with ribosomal profiling during early (left) and late (right) exponential growth phases^12^. (b) Time profiles are shown for viable cell density (VCD) and titer in experimental culture. Shaded boxes indicate the time points corresponding to early (day 3) and late (day 6) growth phases. (c) Flux balance analysis was used to predict specific productivity (q_p_) with the iCHO2048s model before and after in silico knockout of *neoR* gene. (d) Growth-productivity trade-offs were predicted by iCHO2048s and demonstrated a potential 18% increase after the *neoR* in silico knockout. The formula for calculating the trade-off improvement (*Δ*) is shown in the plot. L_WT_ = length of trade-off curve before knockout, L_KO_ = length of trade-off curve after knockout. (e) Ribosomal occupancy for all mRNA sequences bearing a signal peptide sequence were analyzed from the Kallehauge et al.^12^ study and demonstrated that the top 30 secreted proteins accounted >50% of the ribosomal occupancy of secreted proteins. Error bars represent the standard deviation of three biological replicates. Source data are provided as a Source Data file.

## Discussion

Mammalian cells synthesize and process thousands of proteins through their secretory pathway. Many of these proteins, including hormones, enzymes, and receptors, are essential for mediating mammalian cell interactions with their environment. Therefore, many have therapeutic importance either as drugs or as targets. The expression and secretion of recombinant proteins represents a significant anabolic demand that drains several substrates from cellular metabolism (e.g., amino acids, sugar nucleotides, ATP)^21,22^. Furthermore, the recombinant proteins demand adequate expression of supporting proteins involved in their transcription, translation, folding, modification, and secretion. Thus, there has been an increasing interest in engineering the mammalian secretory pathway to boost protein production^23–26^. Despite important advances in the field^27^, current strategies to engineer the secretory pathway have remained predominantly empirical^28,29^. Recent modeling approaches, however, have enabled the analysis of the metabolic capabilities of important eukaryotic cells under different genetic and environmental conditions^17,30–32^. With the development of genome-scale models of protein-producing cells, such as CHO^17^, HEK-293^33^ and hybridomas^34,35^, it is now possible to gain a systems-level understanding of the mammalian protein production phenotype^36^.

Efforts have been underway to enumerate the machinery needed for protein production. For example, Lund and colleagues^6^ recently reconstructed a comprehensive genetic network of the mouse secretory pathway. By comparing the mouse and CHO-K1 genomes and mapping CHO gene expression data onto this network, the authors identified potential targets for CHO cell engineering, demonstrating the potential of systems biology to interrogate and understand protein secretion in animal cells. This genetic network reconstruction, although useful for contextualizing omics data (e.g., RNA-Seq), is not set up for simulations of protein production, nor integrated with additional cellular processes such as metabolism. Therefore, our work is complementary in that it allows one to also to quantify the cost and cellular capacity for protein production by delineating the mechanisms of all biosynthetic steps and bioenergetic processes in the cell.

Here we presented the first genome-scale reconstruction of the secretory pathway in mammalian cells coupled to metabolism. We connected this to current metabolic networks, yielding models of protein secretion and metabolism for human, mouse and CHO cells. These models compile decades of research in biochemistry and cell biology of higher eukaryotes and present it in a mathematical model. Using our model, we quantitatively estimated the energetic cost of producing several therapeutic proteins and all proteins in the CHO cell and human secretomes. We also identified factors limiting the secretion of individual products and observed that these depend on both the complexity of the product and the composition of the culture media. Furthermore, by integrating ribosomal profiling data with our model we found that CHO cells have selectively suppressed the expression of energetically expensive secreted proteins. Expanding upon this observation, we demonstrated that specific productivities can be predictably increased following the knock-down of an energetically expensive, non-essential protein. Furthermore, consistent with this, we have recently shown more than 50% reductions in total host cell protein production, along with increases in mAb titer when deleting 14 highly abundant proteins in CHO cells. Further studies will likely further explore how much of the CHO cell proteome can be deleted to further enhance recombinant protein secretion^20^.

It is important to note that while our models capture major features of secreted proteins, there are additional PTMs (e.g., phosphorylation, gamma carboxylation), pathway machinery (e.g., chaperones), and cell processes that could possibly be captured in further expansions of the modeling framework^6^ (e.g., the unfolded protein response). These could be included as energetic costs associated with building and maintaining the secretory machinery (chaperones^3^, disulfide oxidoreductases^37^, glycosyltransferases^38^); protein stability and turnover rates^39^; solubility constraints^40^ and molecular crowding effects^41^. As these are captured by the models in a protein product-specific manner, predictions of protein production capacity will improve, and the models could provide further insights for cell engineering for biotechnology or to obtain a deeper understanding of mechanisms underlying amyloid diseases. Finally, a simplification of our secretory model is that it only computes the bioenergetic cost of synthesizing and attaching single representative N- and O-linked glycans to secreted proteins (i.e., it does not include the microheterogeneity and diversity of glycan structures of different proteins). Thus, an immediate potential expansion of our secretory model would involve coupling it to existing computational models of protein glycosylation^42,43^. For example, given an N-glycan reaction network that captures the glycoform complexity of a target protein^44^, one could build secretory reactions for the specific glycoforms of interest and compute the metabolic demands associated with each of them to identify potential targets and nutrient supplementations for glycoengineering.

In conclusion, the results of our study have important implications regarding the ability to predict protein expression based on protein specific attributes and energetic requirements. The secretory pathway models here stand as novel tools to study mammalian cells and the energetic trade-off between growth and protein secretion in a product- and cell-specific manner. We presented algorithms that provide novel insights with our models, and expect that many other methods can be developed to answer a wide array of questions surrounding the secretory pathway, as seen for metabolism^45^. To facilitate further use of these models, we provide our code and detailed instructions on how to construct protein-specific models in the Jupyter Notebooks available at https://github.com/LewisLabUCSD/MammalianSecretoryRecon.

## Methods

### Reconstruction of the mammalian secretory pathway

A list of proteins and enzymes in the mammalian secretory pathway was compiled from literature curation, UniProt, NCBI Gene, NCBI Protein and CHOgenome.org (see Supplementary Data 1). To facilitate the reconstruction process, the secretory pathway was divided into twelve subsystems or functional modules (Fig. 1) to sort the components according to their function. These subsystems correspond to the major steps required to process and secrete a protein. The components from a prior yeast secretory pathway reconstruction^3^ were used as a starting reference. To build species-specific models, orthologs for human, mouse and the Chinese hamster were identified and used, while yeast components and subsystems that are not present in the mammalian secretory pathway were removed. Additional subsystems were added when unique to higher eukaryotes, such as the calnexin-calreticulin cycle in the ER^46^. These were constructed de novo and added to the reconstruction. The databases and literature were then consulted to identify the remaining components involved in each subsystem of the mammalian secretory pathway. Since most components in the mammalian secretory pathway have been identified in mouse and human, BLAST was utilized to identify the corresponding Chinese hamster orthologs by setting human as the reference organism and a cutoff of 60% of sequence identity. See Supplementary Discussion for an overview of the mammalian secretory pathway and its comparison with the yeast secretory pathway.

### Protein Specific Information Matrix (PSIM)

The PSIM (Supplementary Data 2) contains the necessary information to construct a protein-specific secretory model from the template reactions in our reconstruction. The columns in the PSIM are presence of a signal peptide (SP), number of disulfide bonds (DSB), presence of Glycosylphosphatidylinositol (GPI) anchors, number of N-linked (NG) and O-linked (OG) glycans, number of transmembrane domains (TMD), subcellular location, protein length, and molecular weight. For most proteins, the information in the PSIM was obtained from the Uniprot database. When necessary, computational tools were used to predict signal peptides (PrediSi^7^) and GPI anchors (GPI-SOM^8^). Finally, additional information on the number of O-linked glycosylation sites of certain proteins were obtained from experimental data in previous studies^9,47^. The PSIMs of the CHO and human secretomes are a subset of the full PSIM and contains only the proteins with a signal peptide (predicted or confirmed in Uniprot). The distribution of all PTMs across the human, mouse and CHO proteomes can be found in Jupyter Notebook D. For analyzing secretomes, a total of 3378 human proteins were picked based on the presence of a signal peptide in their sequence according to their annotation in the UniProt database. Similarly, 5,641 CHO proteins were picked based on the presence of a signal peptide in their sequence and/or for being localized in the cell membrane according to the UniProt database.

### Detection of N-linked glycosylation sites in CHO proteome

The number of N-linked glycosylation sites in the PSIM was determined computationally and experimentally as follows. CHO-K1 cells (ATCC) were lysed, denatured, reduced, alkylated and digested by trypsin. Desalted peptides were incubated with 10 mM sodium periodate in dark for 1 hour before coupling to 50 μL of (50% slurry) hydrazide resins. After incubation overnight, non-glycosylated peptides were washed with 1.5 M NaCl and water. The N-glycosylated peptides were released with PNGaseF at 37 °C and desalted by using a C18 SepPak column. Strong cation exchange (SCX) chromatography was used to separate the sample into 8 fractions. Each fraction was analyzed on an LTQ-Orbitrap Velos (Thermo Electron, Bremen, Germany) mass spectrometer. During the mass spectrometry data analysis, carbamidomethylation was set as a fixed modification while oxidation, pyroglutamine and deamidation were variable modifications.

### Construction of models and constraint-based analysis

We wrote a Jupyter Notebook in Python (see Jupyter Notebook A) that takes a row from the PSIM as input to produce an expanded iCHO2048s, Recon 2.2s, or iMM1685s metabolic model with the product-specific secretory pathway of the corresponding protein. Flux balance analysis (FBA^48^) and all other constraint-based analyses were done using the COBRA toolbox v2.0^49^ in MATLAB R2015b and the Gurobi solver version 6.0.0. The analyses in Figs. 2, 3, and 4 were done using the constraints in the Supplementary Data 3. For the iCHO2048s models secreting human proteins, we set the same constraints in all models and computed the theoretical maximum productivity (*max*_*q*p_) while maintaining a growth rate (in units of inverse hours) of 0.01. Finally, since the exact glycoprofiles of most proteins in CHO are unknown and some even change over time in culture^50^, we simplified our models by only adding the core N-linked and O-linked glycans to the secreted proteins.

### Batch cultivation

Two isogenic CHO-S cell lines (Thermo Fisher Scientific, USA) adapted to grow in suspension, one producing Enbrel (Etanercept) and the other producing human plasma protease C1 inhibitor (C1INH), were seeded at 3 × 10^5^ cells per mL in 60 mL CD-CHO medium (Thermo Fisher Scientific, USA) supplemented with 8 mM L-Glutamine (Lonza) and 1 μL per mL anti-clumping agent (Life Technologies), in 250 mL Erlenmeyer shake flasks. Cells were incubated in a humidified incubator at 37°C, 5% CO_2_ at 120 rpm. Viable cell density and viability were monitored every 24 hours for 7 days using the NucleoCounter NC-200 Cell Counter (ChemoMetec). Daily samples of spent media were taken for extracellular metabolite concentration and titer measurements by drawing 0.8 mL from each culture, centrifuging it at 1000 g for 10 minutes and collecting the supernatant and discarding the cell pellet.

### Titer determination

To quantify Enbrel and C1INH titers, biolayer interferometry was performed using an Octet RED96 (Pall Corporation, Menlo Park, CA). ProA biosensors (Fortebio 18-5013) were hydrated in PBS and preconditioned in 10 mM glycine pH 1.7. A calibration curve was prepared using Enbrel (Pfizer) or C1INH at 200, 100, 50, 25, 12.5, 6.25, 3.13, 1.56, 0.78 μg per ml. Culture spent media samples were collected after centrifugation and association was performed for 120 s with a shaking speed of 200 rpm at 30 °C. Octet System Data Analysis 7.1 software was used to calculate binding rates and absolute protein concentrations.

### Extracellular metabolite concentration measurements

The concentrations of glucose, lactate, ammonium (NH_4_^+^), and glutamine in spent media were measured using the BioProfile 400 (Nova Biomedical). Amino acid concentrations were determined via High Performance Liquid Chromatography using the Dionex Ultimate 3000 autosampler at a flow rate of 1mL per minute. Briefly, samples were diluted 10 times using 20 μL of sample, 80 μL MiliQ water, and 100 μL of an internal amino acid standard. Derivatized amino acids were monitored using a fluorescence detector. OPA-derivatized amino acids were detected at 340ex and 450em nm and FMOC-derivatized amino acids at 266ex and 305em nm. Quantifications were based on standard curves derived from dilutions of a mixed amino acid standard (250 ug per mL). The upper and lower limits of quantification were 100 and 0.5 ug permL, respectively.

### Estimation of protein secretion cost

We estimated the energetic cost of synthesizing and secreting all 5,641 endogenous CHO cell proteins and 3,538 endogenous human proteins. These proteins were chosen for containing a signal peptide in their sequence and/or for being localized in the cell membrane (according to the UniProt database). The energetic cost (in units of number of ATP equivalents) of secreting each protein (length L) was computed using the following formulas and assumptions.

#### Energy cost of translation

For each protein molecule produced, 2L ATP molecules are cleaved to AMP during charging of the tRNA with a specific amino acid; 1 GTP molecule is consumed during initiation and 1 GTP molecule for termination; L-1 GTP molecules are required for the formation of L-1 peptide bonds; L-1 GTP molecules are necessary for L-1 ribosomal translocation steps. Thus, the total cost of translation (assuming no proofreading) is 4L.

#### Average cost of signal peptide degradation

On average, signal peptides have a length of 22 amino acids. Thus, the average cost of degrading all peptide bonds in the signal peptide is 22. This average cost was assigned to all proteins analyzed.

#### Energetic cost of translocation across the ER membrane

During activation of the translocon, 2 cytosolic GTP molecules are hydrolyzed. From there, a GTP molecule bound to the folding-assisting chaperone BiP is hydrolyzed to GDP for every 40 amino acids that pass through the translocon pore^46^. Thus, the cost of translocation is (L÷40) + 2.

#### Energetic cost of vesicular transport and secretion

We used published data^51–53^ (see Supplementary Data 1) to compute stoichiometric coefficients for reactions involving vesicular transport. That is, the number of GTP molecules bound to RAB and coat proteins in each type of vesicle (COPII and secretory vesicles). We found that a total of 192 and 44 GTPs must be hydrolyzed to transport one COPII or secretory (i.e. clathrin coated) vesicle from the origin membrane to the target membrane, respectively. Since vesicles do not transport a single protein molecule at a time, we estimated the number of secreted protein molecules that would fit inside a spherical vesicle (see estimated and assumed diameters in the Supplementary Data 1). For that, we assumed that the secreted protein is globular and has a volume V_P_ (nm^3^) that is directly proportional to its molecular weight MW^54^:

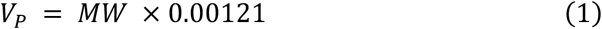

Finally, we assumed that only 70 percent of the vesicular volume can be occupied by the target protein. Thus, the cost of vesicular transport via COPII vesicles with Volume V_COPII_ is:

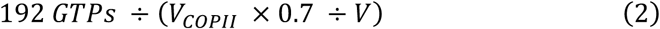

Similarly, the cost of vesicular secretion is:

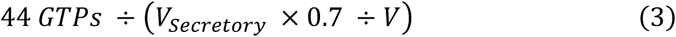

### Constraints used in models and Pareto optimality frontiers

All models were constrained using different sets of experimental uptake rates, which can be found in Supplementary Data 3. To construct Pareto optimality frontiers, we used the robustAnalysis function from the COBRA Toolbox v2.0 in Matlab 2015b using biomass as the control and secretion of the recombinant protein as the objective reactions, respectively.

### Analysis of gene expression versus protein cost

Ribosome-profiling data^12^ were used to quantify the ribosomal occupancy of each transcript in CHO cells. A cutoff of 1 RPKM was used to remove genes with low expression (10,045 genes removed from day 3 analysis and 10,411 from day 6 analysis). We used Spearman correlation to assess the variation of expression levels with respect to protein ATP cost.

### CHO-DG44 model and prediction of *neoR* knock-out effect

Ribosome-profiling data, specific productivity, product sequence, and growth rates of an IgG-producing CHO-DG44 cell line were obtained from a previous publication^12^. From the same cultures, we obtained further cell dry weight and metabolomic data from spent culture medium for this study. The mCADRE algorithm^55,56^ was used to construct a DG44 cell line-specific iCHO2048s model. The specific productivity and the RPKM values of the secreted IgG were used to estimate the translation rate for the *neoR* selection marker gene. We assumed that the flux (in units of mmol per gram dry weight per hour) through the *neoR* translation reaction (*v*_neoR_) should be proportional to that of the IgG translation rate (*v*_IgG_, calculated from the measured specific productivity) and related to their expression ratios (i.e. the RPKM values of their genes in the ribosome-profiling data).

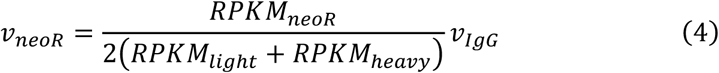

Finally, a reaction of *neoR* peptide translation (which is expressed in cytosol and is not processed in the secretory pathway) was added to construct a *neoR*-specific iCHO2048s model. Uptake and secretion rates of relevant metabolites on days 3 and 6 of cell culture were used to constrain our model. Because recombinant proteins represent 20% of total cell protein^57^, we scaled the coefficients of all 20 amino acids in the model’s biomass reaction accordingly (i.e. each coefficient was multiplied by 0.8). We then used FBA to predict the specific productivity of IgG with or without *neoR*.

### Cell dry weight measurements

For cell dry weight measurements, 6 tubes containing 2 mL of culture samples of known viable cell density and viability were freeze dried, weighed, washed in PBS, and weighed again. The difference in weight was used to calculate the mass per cell. The procedure resulted in an average cell dry weight of 456 pg per cell. As a simplification, we assumed that cell dry weight does not significantly differ from this average measured value during culture and thus was used when computing flux distributions in all simulations.

### Calculation of growth and productivity rates

Supplementary Data 3 contains the experimental uptake and secretion rates used to constrain the iCHO2048s models^12,22,23^. When rates were not explicitly stated in the studies we consulted, we used a method we developed previously^27^. Briefly, appropriate viable cell density, titer, and metabolite concentration plots were digitized using WebPlot Digitizer software and we computed the corresponding rates as follows:

Growth rate (in units of inverse hours):

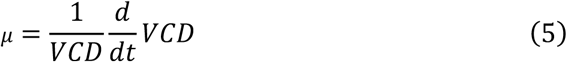

Where VCD is the viable cell density (in units of cells per milliliter)

Specific productivity (in units of picograms per cell per hour):

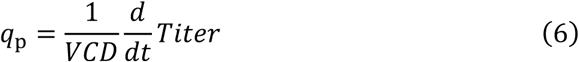

Consumption or production rate *v*_x_ of metabolite x (in units of millimoles per gram dry weight per hour):

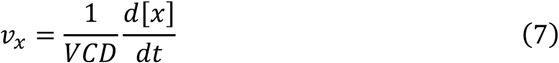

## Data Availability

All data that support the findings of this study, including the models, tables, and Jupyter Notebooks, are available at https://github.com/LewisLabUCSD/MammalianSecretoryRecon as well as in the Supplementary Data and Source Data files. The Ribo-seq and RNA-seq data from the study by Kallehauge et al.^12^ is available on the Gene Expression Omnibus with GEO Accession Number GSE79512 (https://www.ncbi.nlm.nih.gov/geo/query/acc.cgi?acc=GSE79512). The RNA sequencing data for human tissue is freely available at the Human Protein Atlas website (https://www.proteinatlas.org/about/download).

## Code Availability

All code used to generate the results of this study, including Jupyter Notebooks, MATLAB, and Python scripts, are freely accessible at https://github.com/LewisLabUCSD/MammalianSecretoryRecon

## Acknowledgements

The authors would like to thank Philipp Spahn, Austin Chiang, and Chih-Chung Kuo for their insightful comments on this manuscript. This work was supported by generous funding from the Novo Nordisk Foundation provided to the Center for Biosustainability at the Technical University of Denmark (NNF10CC1016517 and NNF16CC0021858), and from NIGMS (R35 GM119850), and a fellowship from the Government of Mexico (CONACYT) and the University of California Institute for Mexico and the United States (UC-MEXUS).

## Author Contributions

J.M.G., A.F., and N.E.L. conceived the project and designed the experiments. J.M.G., A.F., S.L., T.B.K, H.H., L.G., D.L collected data. J.M.G. and D.B.H. performed experiments. J.M.G. performed research, analyzed the data and interpreted the results. J.M.G., A.F. and N.E.L. drafted the article. M.J.B., B.V., H.F.K., G.M.L., B.O.P, J.N., and N.E.L. critically revised and approved the article.

## Supplementary Information

**Supplementary Figure 1.**
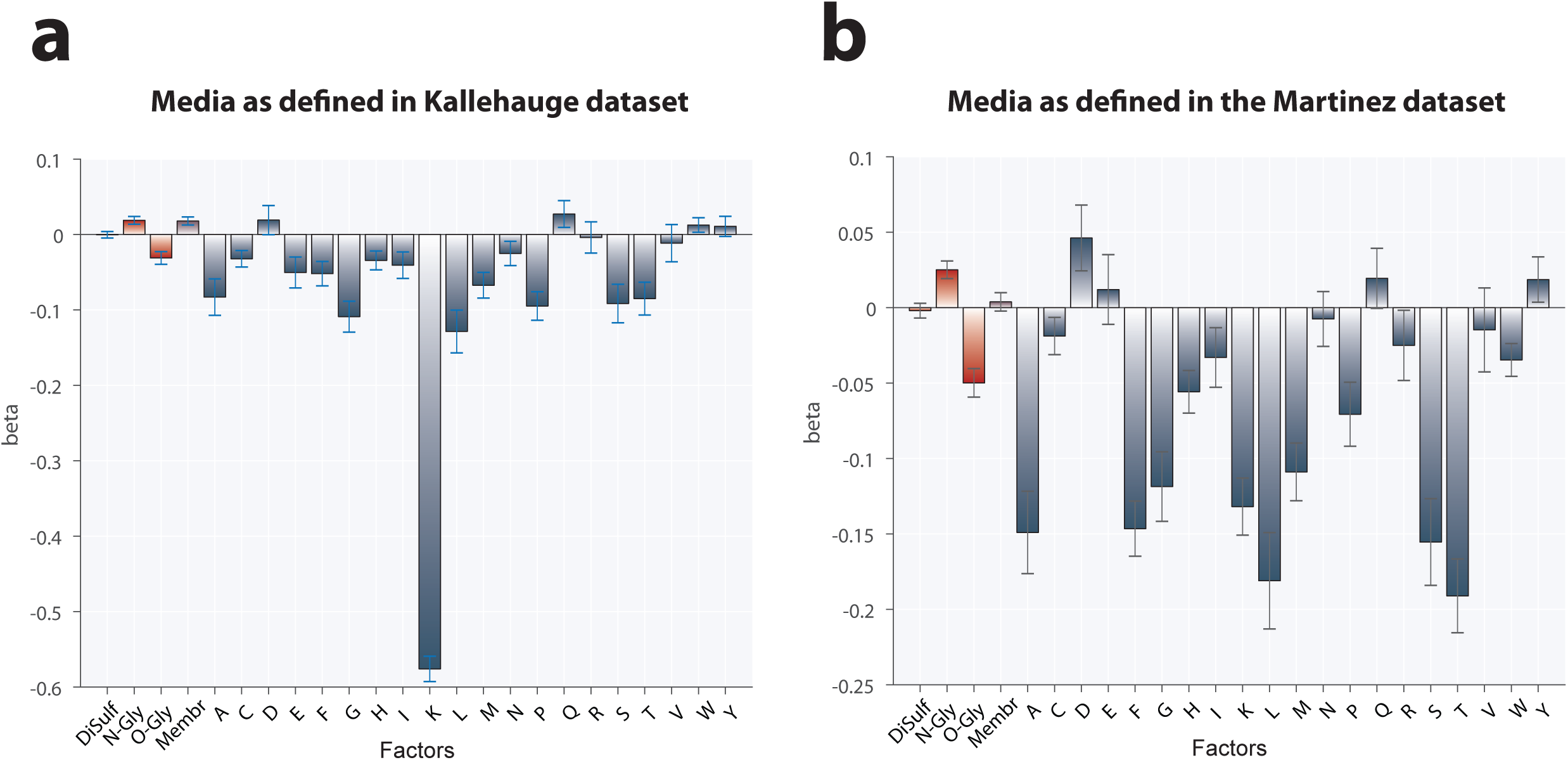
Factors affecting iCHO2048s-predicted productivity with two different media compositions. Linear regression coefficients (β) to quantify the contribution of PTMs to the explained variation in specific productivity using uptake rates different from those used in Figure 4c. The specific consumption rates are listed in Supplementary Table 3 as Kallehauge^12^ (left panel) and Martinez^59^ (right panel). Error bars represent the standard error of the fitted coefficients. Source data are provided as a Source Data file.

**Supplementary Figure 2.**
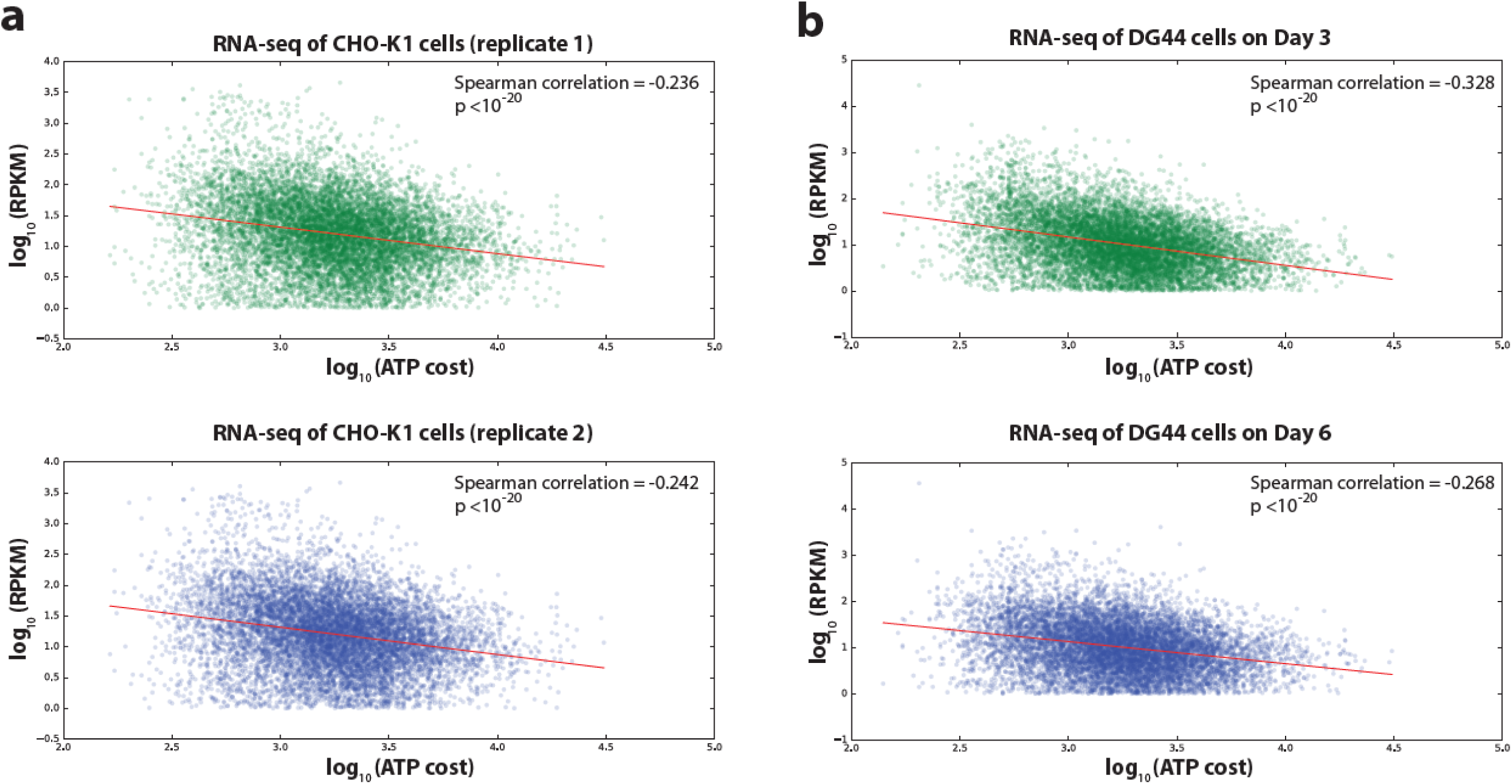
Spearman correlation between ATP cost and gene expression levels in non-producing CHO-K1 and CHO-DG44 mAb-producing cells. Gene transcription levels from (a) van Wijk et al.^13^ and (b) Kallehauge et al.^12^ were compared against the ATP cost of producing the translated proteins. Source data are available as a Source Data file.

**Supplementary Figure 3.**
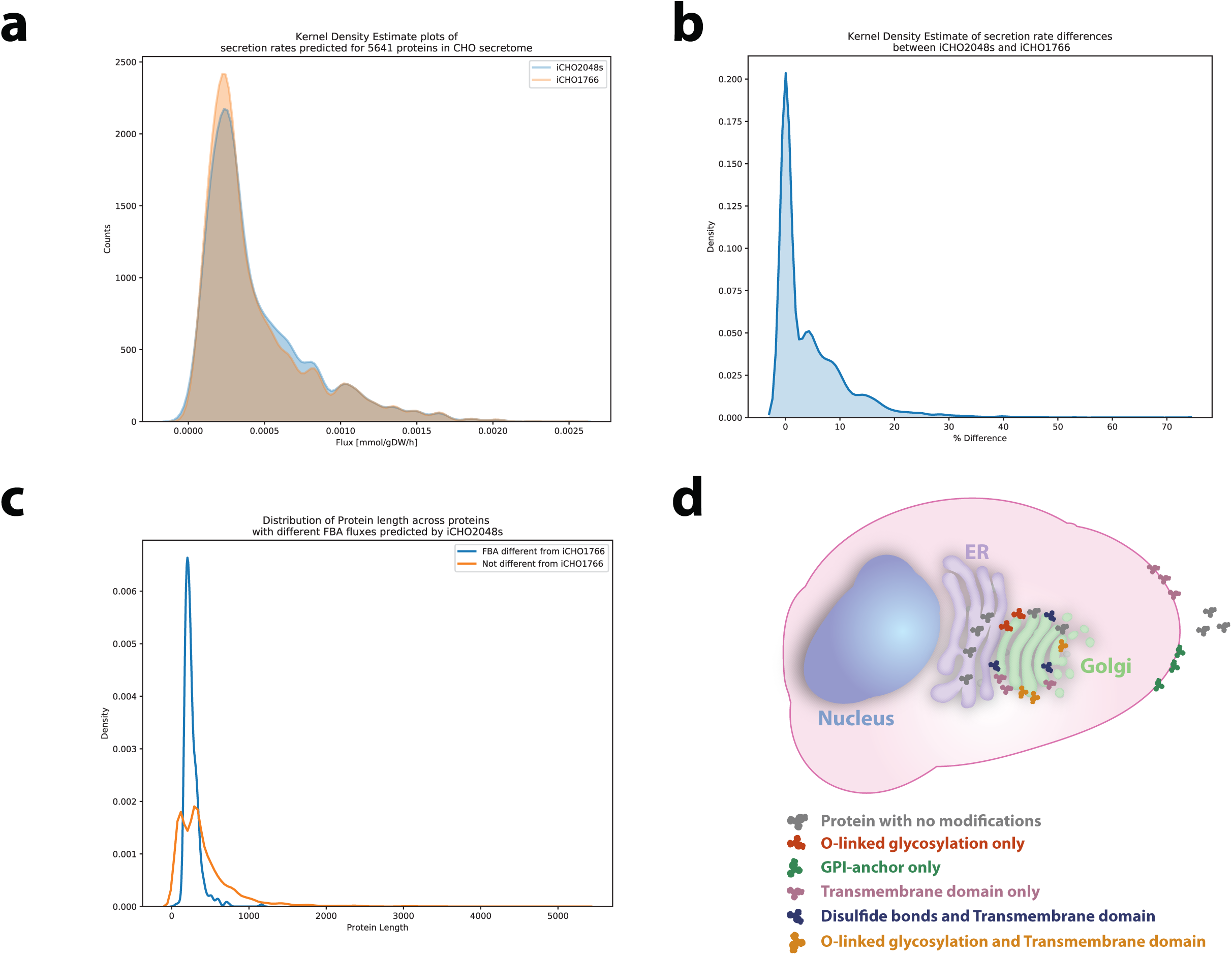
Comparison of secretion rates predicted by iCHO2048s and iCHO1766. Kernel Density Plots of (a) secretion rates for 5641 proteins in the CHO secretome, as computed with iCHO2048s (blue) and iCHO1766 (red), (b) the percentage difference between predictions with iCHO2048s and iCHO1766, and (c) the protein lengths (in units of amino acids in sequence) of proteins showing a secretion rate difference in both models (blue) or not (red). iCHO2048s predicts different fluxes for proteins with a specific post-translational modification profile, size, and localization. For about 8% of the target secretome, secretion rates predicted with iCHO2048s are at least 15% different from their iCHO1766 counterparts. Interestingly, this 8% corresponds to short (less than 350 amino acids) secreted proteins with O-linked glycans, GPI anchors or transmembrane domains whose final location is the extracellular space, the ER lumen, the Golgi membrane, or the plasma membrane, as summarized in (d). Thus, for a proportion of the secretome, there are non-negligible energetic and synthetic costs associated with vesicular transport, protein folding, and membrane anchoring only accounted for when iCHO2048s couples to metabolism. A detailed description of the results, as well as the source data, can be found in Jupyter Notebook E (https://github.com/LewisLabUCSD/MammalianSecretoryRecon/JUPYTER_NOTEBOOKS).

## Supplementary Discussion

### Overview of the Secretory Pathway in animal cells

Historically, most of the knowledge on the secretory pathway was obtained by studying protein transport processes and secretion in *Saccharomyces cerevisiae*^1^. Albeit quite similar in core functions, the secretory pathways of mammalian cells and fungi differ significantly in some of the steps which have been evolved based on species-specific secretion phenotypes^2^. The following paragraphs briefly overview the mammalian secretory pathway and highlights pathways exclusive to animals not present in fungi. The last section provides an in-depth comparison of the yeast and animal secretory pathways while highlighting the most important differences between both.

#### Translocation and processing in endoplasmic reticulum

Proteins destined to the secretory pathway generally bear a signal peptide at the amino-terminus which targets the proteins to the endoplasmic reticulum (ER) where the initial post-translational modifications (PTMs) take place. This transport requires translocating the target protein across the ER membrane through two general pathways: co-translational translocation (GTP dependent) and post-translational translocation (ATP dependent)^3^. An additional pathway for tail-anchored (TA) proteins into the ER membrane has also been discussed in the literature and included in our iCHO1921s reconstruction^4,5^. Once inside the ER lumen, the target proteins are folded by the action of several transmembrane ER proteins, including calnexin, calreticulin, and other luminal chaperones^6–8^. In the event of protein misfolding, a target protein may go through a “quality control” system (exclusive in the mammalian secretory pathway) that attempts to correct for folding errors^9,10^. However, if the misfolded state of the protein is sustained for too long, the protein then enters the ER associated degradation pathway, or ERAD, which involves retrotranslocation of the misfolded protein back to the cytosol, ubiquitination and proteasomal degradation^11–13^.

Besides folding, a target protein may acquire additional PTMs while inside the ER such as attachment of a glycosylphosphatidylinositol (GPI) anchor^14,15^, formation of disulfide bonds^16^, and N-linked glycosylation^17–20^. After these PTMs are successfully completed, the target proteins are transported to the Golgi apparatus via COPII-coated vesicles that bud from the ER^21,22^ whereas misfolded proteins are retro-translocated to the cytoplasm^23,24^ for proteasomal degradation via the ER-associated degradation pathway (ERAD)^25,26^. In the Golgi apparatus, N-glycans are processed into branched and complex glycoforms and proteins are further glycosylated with O-linked glycans^27–29^ and then sorted to their final destination (e.g. lysosome, extracellular medium) via clathrin-coated secretory vesicles^30–33^.

#### A note on translocation pathways

In co-translational translocation, proteins destined to the secretory pathway bear a hydrophobic signal sequence at the amino-terminus that promotes the targeting of ribosome-nascent chain (RNC) complexes to the ER via binding to the signal recognition particle (SRP). The SRP recognizes the signal peptide as soon as it emerges from the ribosome during translation. Then, the newly formed SRP-RNC complex is recognized by the SRP receptor on the ER membrane where translocation is initiated by interaction with the Sec61 complex (Sec61C) and assisted by the chaperone BiP to increase the efficiency and ensure the unidirectionality of this process^30^.

Post-translational translocation, in contrast to co-translational translocation, occurs independently of SRP and its receptor^34^. Furthermore, this process does not rely too heavily on the Sec61C to translocate the target protein and instead utilizes the protein Sec62 as a safe route that guarantees the efficient translocation of small proteins (<160 amino acids in length)^35^.

Finally, the pathway for inserting TA proteins into the ER membrane also occurs post-translationally due to the fact that the ER targeting signal of TA proteins is located very close to the carboxy-terminus, which allows the ribosome to release the protein before it is recognized and localized to the ER^36^. This pathway depends on ATP and one of the main players in the process is a transmembrane recognition complex known as TRC40 or Asna1^37^.

## Important differences between the yeast and animal secretory pathways

As mentioned above, core functions of the secretory pathway are conserved between mammalian and yeast cells. These core functions (see Table SD.2) are:

- Translocation through endoplasmic reticulum
- Primary glycosylation in ER (N-linked glycans) and Golgi (N-linked and O-linked glycans)
- Protein folding and quality control in ER
- Anterograde and retrograde vesicular transport between ER and Golgi via COPII and COPI vesicles, respectively.
- Dolichol pathway for N-linked core glycan translocation through the ER membrane
- Endoplasmic reticulum associated degradation (ERAD)
- GPI biosynthesis
- Unfolded protein response (UPR)

Nevertheless, minor and major differences exist between the yeast and mammalian secretory pathways. Some of these differences have been thoroughly reviewed before in an excellent review by Delic and colleagues^2^ and are summarized in Table SD.1 below. Here, we highlight the major differences between both secretory pathways that are relevant for modeling purposes using the secretory reconstructions.

**Table SD.1.** Summary of differences between mammalian and yeast secretory pathways as described by Delic et al.^2^.

| Description of difference | Mammalian secretory pathway | Yeast secretory pathway | Importance for modeling purposes |
| --- | --- | --- | --- |
| Chaperones involved in translocation | The main chaperone is BiP | The main chaperone is Kar2 | Minor |
| Presence of heat-shock proteins (HSPs) in ER | Mainly presence of proteins in the Hsp90 family | Not found in yeast | Minor |
| Enzymes for detoxification of reactive oxygen species in ER | Contains several enzymes such as Ero1 and glutathione peroxidases | Not found in yeast | Major |
| Oxidation state of Protein disulfide isomerase (PDI) | PDI is mainly reduced | PDI is mainly oxidized | Minor |
| Components of calnexin-calreticulin cycle | Includes an enzyme coded by the UGGT gene to transfer glucose residues to core N-linked glycans in misfolded proteins | Lacks UGGT and instead directs misfolded proteins to ER exit | Major |
| ERAD pathway branches for degrading misfolded proteins | Capable of directing misfolded proteins towards the ERAD pathway by trimming N-linked glycan residues in the A, B and C branches | Capable of directing misfolded proteins towards the ERAD pathway by trimming N-linked glycan residues only in B and C branches | Major |
| Components of COPII vesicles | Contains four isoforms of Sec24 | Expresses Sec24 with three cargo binding sites as well as Sec24 homologs Sfb2-3 | Minor |

Finally, the table below summarizes the differences between the mammalian and the fungal secretory pathway reconstructions in terms of components, reactions, and subsystems.

**Table SD.2.** Overview of main differences between the mammalian and yeast secretory pathway reconstructions.

| Secretory pathway reconstruction | Number of components | Number of reactions | Number of Subsystems | Core subsystems (in both mammalian and yeast secretory pathways) | Unique subsystems |
| --- | --- | --- | --- | --- | --- |
| Mammalian | 271 | 144 | 12 | A total of 9 core subsystems: COPI, COPII, Dolichol pathway, ER glycosylation, ERAD, Golgi processing, GPI biosynthesis, Protein folding, and Translocation | Clathrin vesicles, GPI transfer |
| Yeast | 165 | 137 | 16 |  | ALP pathway, CPY pathway |

## References

1. Uhlén, M. et al. Tissue-based map of the human proteome. Science 347, (2015).

2. Walsh, G. Biopharmaceutical benchmarks 2018. Nat. Biotechnol. 36, 1136–1145 (2018).

3. Feizi, A., Österlund, T., Petranovic, D., Bordel, S. & Nielsen, J. Genome-Scale Modeling of the Protein Secretory Machinery in Yeast. PLoS One 8, e63284 (2013).

4. Liu, L., Feizi, A., Österlund, T., Hjort, C. & Nielsen, J. Genome-scale analysis of the high-efficient protein secretion system of Aspergillus oryzae. BMC Syst. Biol. 8, 73 (2014).

5. Irani, Z. A., Kerkhoven, E. J., Shojaosadati, S. A. & Nielsen, J. Genome-scale metabolic model of Pichia pastoris with native and humanized glycosylation of recombinant proteins. Biotechnol. Bioeng. 113, 961–969 (2016).

6. Lund, A. M. et al. Network reconstruction of the mouse secretory pathway applied on CHO cell transcriptome data. BMC Syst. Biol. 11, 37 (2017).

7. Hiller, K., Grote, A., Scheer, M., Munch, R. & Jahn, D. PrediSi: prediction of signal peptides and their cleavage positions. Nucleic Acids Res. 32, W375–W379 (2004).

8. Fankhauser, N. & Maser, P. Identification of GPI anchor attachment signals by a Kohonen self-organizing map. Bioinformatics 21, 1846–1852 (2005).

9. Yang, Z. et al. The GalNAc-type O-Glycoproteome of CHO cells characterized by the SimpleCell strategy. Mol. Cell. Proteomics 13, 3224–35 (2014).

10. Kaufman, R. J. et al. Effect of von Willebrand Factor Coexpression on the Synthesis and Secretion of Factor VIII in Chinese Hamster Ovary Cells. Mol. Cell. Biol. 9, 1233–1242 (1989).

11. Pipe, S. W., Morris, J. A., Shah, J. & Kaufman, R. J. Differential interaction of coagulation factor VIII and factor V with protein chaperones calnexin and calreticulin. J. Biol. Chem. 273, 8537–44 (1998).

12. Kallehauge, T. B. et al. Ribosome profiling-guided depletion of an mRNA increases cell growth rate and protein secretion. Sci. Rep. 7, 40388 (2017).

13. van Wijk, X. M. et al. Whole-Genome Sequencing of Invasion-Resistant Cells Identifies Laminin α2 as a Host Factor for Bacterial Invasion. MBio 8, e02128–16 (2017).

14. Feizi, A., Gatto, F., Uhlen, M. & Nielsen, J. Human protein secretory pathway genes are expressed in a tissue-specific pattern to match processing demands of the secretome. npj Syst. Biol. Appl. 3, 22 (2017).

15. Swainston, N. et al. Recon 2.2: from reconstruction to model of human metabolism. Metabolomics 12, 109 (2016).

16. Sigurdsson, M. I., Jamshidi, N., Steingrimsson, E., Thiele, I. & Palsson, B. O. A detailed genome-wide reconstruction of mouse metabolism based on human Recon 1. BMC Syst. Biol. 4, 140 (2010).

17. Hefzi, H. et al. A Consensus Genome-scale Reconstruction of Chinese Hamster Ovary Cell Metabolism. Cell Syst. 3, 434–443.e8 (2016).

18. Templeton, N., Dean, J., Reddy, P. & Young, J. D. Peak Antibody Production is Associated With Increased Oxidative Metabolism in an Industrially Relevant Fed-Batch CHO Cell Culture. Biotechnol. Bioeng 110, 2013–2024 (2013).

19. Kim, S. J., Kim, N. S., Ryu, C. J., Hong, H. J. & Lee, G. M. Characterization of chimeric antibody producing CHO cells in the course of dihydrofolate reductase-mediated gene amplification and their stability in the absence of selective pressure. Biotechnol. Bioeng. 58, 73–84 (1998).

20. Kol, S. et al. Multiplex secretome engineering enhances recombinant protein production and purity. Preprint at https://www.biorxiv.org/content/10.1101/647214v1 (2019).

21. Gu, M. B., Todd, P. & Kompala, D. S. Metabolic burden in recombinant CHO cells: effect of dhfr gene amplification and lacZ expression. Cytotechnology 18, 159–166 (1996).

22. Gu, M. B., Todd, P. & Kompala, D. S. Analysis of foreign protein overproduction in recombinant CHO cells. Effect of growth kinetics and cell cycle traverse. Ann. N. Y. Acad. Sci. 721, 194–207 (1994).

23. Hansen, H. G., Pristovšek, N., Kildegaard, H. F. & Lee, G. M. Improving the secretory capacity of Chinese hamster ovary cells by ectopic expression of effector genes: Lessons learned and future directions. Biotechnol. Adv. 35, 64–76 (2017).

24. Delic, M., Göngrich, R., Mattanovich, D. & Gasser, B. Engineering of Protein Folding and Secretion—Strategies to Overcome Bottlenecks for Efficient Production of Recombinant Proteins. Antioxid. Redox Signal. 21, 414–437 (2014).

25. Le Fourn, V., Girod, P.-A., Buceta, M., Regamey, A. & Mermod, N. CHO cell engineering to prevent polypeptide aggregation and improve therapeutic protein secretion. Metab. Eng. 21, 91–102 (2014).

26. Kuo, C. C. et al. The emerging role of systems biology for engineering protein production in CHO cells. Curr. Opin. Biotechnol. 51, 64–69 (2018).

27. Golabgir, A. et al. Quantitative feature extraction from the Chinese hamster ovary bioprocess bibliome using a novel meta-analysis workflow. Biotechnol. Adv. 34, 621–633 (2016).

28. Borth, N., Mattanovich, D., Kunert, R. & Katinger, H. Effect of Increased Expression of Protein Disulfide Isomerase and Heavy Chain Binding Protein on Antibody Secretion in a Recombinant CHO Cell Line. Biotechnol. Prog. 21, 106–111 (2008).

29. Ku, S. C. Y., Ng, D. T. W., Yap, M. G. S. & Chao, S.-H. Effects of overexpression of X-box binding protein 1 on recombinant protein production in Chinese hamster ovary and NS0 myeloma cells. Biotechnol. Bioeng. 99, 155–164 (2008).

30. Yusufi, F. N. K. et al. Mammalian Systems Biotechnology Reveals Global Cellular Adaptations in a Recombinant CHO Cell Line. Cell Syst. 4, 530–542.e6 (2017).

31. Selvarasu, S. et al. Combined in silico modeling and metabolomics analysis to characterize fed-batch CHO cell culture. Biotechnol. Bioeng. 109, 1415–1429 (2012).

32. Gutierrez, J. M. & Lewis, N. E. Optimizing eukaryotic cell hosts for protein production through systems biotechnology and genome-scale modeling. Biotechnol. J. 10, 939–49 (2015).

33. Quek, L.-E. et al. Reducing Recon 2 for steady-state flux analysis of HEK cell culture. J. Biotechnol. 184, 172–178 (2014).

34. Selvarasu, S., Karimi, I. A., Ghim, G.-H. & Lee, D.-Y. Genome-scale modeling and in silico analysis of mouse cell metabolic network. Mol. BioSyst. 6, 152–161 (2009).

35. Sheikh, K., Förster, J. & Nielsen, L. K. Modeling Hybridoma Cell Metabolism Using a Generic Genome-Scale Metabolic Model of Mus musculus. Biotechnol. Prog. 21, 112–121 (2008).

36. Galleguillos, S. N. et al. What can mathematical modelling say about CHO metabolism and protein glycosylation? Comput. Struct. Biotechnol. J. 15, 212–221 (2017).

37. Araki, K. & Inaba, K. Structure, mechanism, and evolution of Ero1 family enzymes. Antioxid. Redox Signal. 16, 790–9 (2012).

38. Jimenez Del Val, I., Polizzi, K. M. & Kontoravdi, C. A theoretical estimate for nucleotide sugar demand towards Chinese Hamster Ovary cellular glycosylation. Sci. Rep. 6, (2016).

39. O’Brien, E. J., Lerman, J. A., Chang, R. L., Hyduke, D. R. & Palsson, B. O. Genome-scale models of metabolism and gene expression extend and refine growth phenotype prediction. Mol. Syst. Biol. 9, 693–693 (2014).

40. Vazquez, A. & Oltvai, Z. N. Macromolecular crowding explains overflow metabolism in cells. Sci. Rep. 6, (2016).

41. Beg, Q. K. et al. Intracellular crowding defines the mode and sequence of substrate uptake by Escherichia coli and constrains its metabolic activity. Proc. Natl. Acad. Sci. U. S. A. 104, 12663–8 (2007).

42. Spahn, P. N. & Lewis, N. E. Systems glycobiology for glycoengineering. Curr. Opin. Biotechnol. 30C, 218–224 (2014).

43. Tejwani, V., Andersen, M. R., Nam, J. H. & Sharfstein, S. T. Glycoengineering in CHO Cells: Advances in Systems Biology. Biotechnol. J. 13, 1700234 (2018).

44. Spahn, P. N. et al. A Markov chain model for N-linked protein glycosylation - towards a low-parameter tool for model-driven glycoengineering. Metab. Eng. 33, 52–66 (2016).

45. Lewis, N. E., Nagarajan, H. & Palsson, B. O. Constraining the metabolic genotype-phenotype relationship using a phylogeny of in silico methods. Nat. Rev. Microbiol. 10, 291–305 (2012).

46. Araki, K. & Nagata, K. Protein Folding and Quality Control in the ER. Cold Spring Harb. Perspect. Biol. 3, a007526–a007526 (2011).

47. Baycin-Hizal, D. et al. Proteomic Analysis of Chinese Hamster Ovary Cells. J. Proteome Res. 11, 5265–5276 (2012).

48. Orth, J. D., Thiele, I. & Palsson, B. Ø. What is flux balance analysis? Nat. Biotechnol. 28, 245–8 (2010).

49. Schellenberger, J. et al. Quantitative prediction of cellular metabolism with constraint-based models: the COBRA Toolbox v2.0. Nat. Protoc. 6, 1290–1307 (2011).

50. Grainger, R. K. & James, D. C. CHO cell line specific prediction and control of recombinant monoclonal antibody *N* -glycosylation. Biotechnol. Bioeng. 110, 2970–2983 (2013).

51. Borner, G. H. H. et al. Multivariate proteomic profiling identifies novel accessory proteins of coated vesicles. J. Cell Biol. 197, 141–160 (2012).

52. Cheng, Y., Boll, W., Kirchhausen, T., Harrison, S. C. & Walz, T. Cryo-electron Tomography of Clathrin-coated Vesicles: Structural Implications for Coat Assembly. J. Mol. Biol. 365, 892–899 (2007).

53. Takamori, S. et al. Molecular Anatomy of a Trafficking Organelle. Cell 127, 831–846 (2006).

54. Liu, J. K. et al. Reconstruction and modeling protein translocation and compartmentalization in Escherichia coli at the genome-scale. BMC Syst. Biol. 8, 110 (2014).

55. Opdam, S. et al. A Systematic Evaluation of Methods for Tailoring Genome-Scale Metabolic Models. Cell Syst. 4, 318–329.e6 (2017).

56. Wang, Y., Eddy, J. A. & Price, N. D. Reconstruction of genome-scale metabolic models for 126 human tissues using mCADRE. BMC Syst. Biol. 6, 153 (2012).

57. González-Leal, I. J. et al. Use of a Plackett-Burman statistical design to determine the effect of selected amino acids on monoclonal antibody production in CHO cells. Biotechnol. Prog. 27, 1709–1717 (2011).

58. Uhlen, M. et al. The human secretome - the proteins secreted from human cells. Preprint at https://www.biorxiv.org/content/10.1101/465815v2 (2018).

59. Martínez, V. S., Buchsteiner, M., Gray, P., Nielsen, L. K. & Quek, L.-E. Dynamic metabolic flux analysis using B-splines to study the effects of temperature shift on CHO cell metabolism. Metab. Eng. Commun. 2, 46–57 (2015).

## Supplementary References

1. Schekman, R. & Novick, P. 23 genes, 23 years later. Cell 116, S13–5, 1 p following S19 (2004).

2. Delic, M. et al. The secretory pathway: exploring yeast diversity. FEMS Microbiol. Rev. 37, 872–914 (2013).

3. Mandon, E. C., Trueman, S. F. & Gilmore, R. Protein translocation across the rough endoplasmic reticulum. Cold Spring Harb. Perspect. Biol. 5, (2013).

4. Shao, S. & Hegde, R. S. Membrane Protein Insertion at the Endoplasmic Reticulum. Annu. Rev. Cell Dev. Biol. 27, 25–56 (2011).

5. Borgese, N. & Fasana, E. Targeting pathways of C-tail-anchored proteins. Biochim. Biophys. Acta - Biomembr. 1808, 937–946 (2011).

6. Gidalevitz, T., Stevens, F. & Argon, Y. Orchestration of secretory protein folding by ER chaperones. Biochim. Biophys. Acta 1833, 2410–24 (2013).

7. Araki, K. & Nagata, K. Protein folding and quality control in the ER. Cold Spring Harb. Perspect. Biol. 3, a007526 (2011).

8. Braakman, I. & Hebert, D. N. Protein folding in the endoplasmic reticulum. Cold Spring Harb. Perspect. Biol. 5, a013201 (2013).

9. Sousa, M. & Parodi, A. J. The molecular basis for the recognition of misfolded glycoproteins by the UDP-Glc:glycoprotein glucosyltransferase. EMBO J. 14, 4196–203 (1995).

10. Caramelo, J. J. & Parodi, A. J. Getting In and Out from Calnexin/Calreticulin Cycles. J. Biol. Chem. 283, 10221–10225 (2008).

11. Xu, C. & Ng, D. T. W. Glycosylation-directed quality control of protein folding. Nat. Rev. Mol. Cell Biol. 16, 742–752 (2015).

12. Xu, C., Wang, S., Thibault, G. & Ng, D. T. W. Futile Protein Folding Cycles in the ER Are Terminated by the Unfolded Protein O-Mannosylation Pathway. Science (80-.). 340, 978–981 (2013).

13. Olzmann, J. A., Kopito, R. R. & Christianson, J. C. The mammalian endoplasmic reticulum-associated degradation system. Cold Spring Harb. Perspect. Biol. 5, (2013).

14. Ikezawa, H. Glycosylphosphatidylinositol (GPI)-anchored proteins. Biol. Pharm. Bull. 25, 409–17 (2002).

15. Fujita, M. & Kinoshita, T. Structural remodeling of GPI anchors during biosynthesis and after attachment to proteins. FEBS Lett. 584, 1670–1677 (2010).

16. Ron, D. & Harding, H. P. Protein-folding homeostasis in the endoplasmic reticulum and nutritional regulation. Cold Spring Harb. Perspect. Biol. 4, (2012).

17. Ron, E. et al. Bypass of glycan-dependent glycoprotein delivery to ERAD by up-regulated EDEM1. Mol. Biol. Cell 22, 3945–3954 (2011).

18. Mohorko, E., Glockshuber, R. & Aebi, M. Oligosaccharyltransferase: the central enzyme of N-linked protein glycosylation. J. Inherit. Metab. Dis. 34, 869–878 (2011).

19. Aebi, M., Bernasconi, R., Clerc, S. & Molinari, M. N-glycan structures: recognition and processing in the ER. Trends Biochem. Sci. 35, 74–82 (2010).

20. Aebi, M. N-linked protein glycosylation in the ER. Biochim. Biophys. Acta - Mol. Cell Res. 1833, 2430–2437 (2013).

21. Lord, C., Ferro-Novick, S. & Miller, E. A. The Highly Conserved COPII Coat Complex Sorts Cargo from the Endoplasmic Reticulum and Targets It to the Golgi. Cold Spring Harb. Perspect. Biol. 5, a013367–a013367 (2013).

22. Dancourt, J. & Barlowe, C. Protein Sorting Receptors in the Early Secretory Pathway. Annu. Rev. Biochem. 79, 777–802 (2010).

23. Bagola, K., Mehnert, M., Jarosch, E. & Sommer, T. Protein dislocation from the ER. Biochim. Biophys. Acta - Biomembr. 1808, 925–936 (2011).

24. Nakatsukasa, K., Brodsky, J. L. & Kamura, T. A stalled retrotranslocation complex reveals physical linkage between substrate recognition and proteasomal degradation during ER-associated degradation. Mol. Biol. Cell 24, 1765–75, S1-8 (2013).

25. Nakatsukasa, K., Kamura, T. & Brodsky, J. L. Recent technical developments in the study of ER-associated degradation. Curr. Opin. Cell Biol. 29, 82–91 (2014).

26. Hebert, D. N., Bernasconi, R. & Molinari, M. ERAD substrates: Which way out? Semin. Cell Dev. Biol. 21, 526–532 (2010).

27. Croset, A. et al. Differences in the glycosylation of recombinant proteins expressed in HEK and CHO cells. J. Biotechnol. 161, 336–348 (2012).

28. Moremen, K. W., Tiemeyer, M. & Nairn, A. V. Vertebrate protein glycosylation: diversity, synthesis and function. Nat. Rev. Mol. Cell Biol. 13, 448–462 (2012).

29. Stanley, P. Golgi glycosylation. Cold Spring Harb. Perspect. Biol. 3, (2011).

30. Bitsikas, V., Corrêa, I. R. & Nichols, B. J. Clathrin-independent pathways do not contribute significantly to endocytic flux. Elife 3, e03970 (2014).

31. Banfield, D. K. Mechanisms of Protein Retention in the Golgi. Cold Spring Harb. Perspect. Biol. 3, a005264–a005264 (2011).

32. Gannon, J., Bergeron, J. J. M. & Nilsson, T. Golgi and Related Vesicle Proteomics: Simplify to Identify. Cold Spring Harb. Perspect. Biol. 3, a005421–a005421 (2011).

33. Ispolatov, I. & Müsch, A. A model for the self-organization of vesicular flux and protein distributions in the Golgi apparatus. PLoS Comput. Biol. 9, e1003125 (2013).

34. Johnson, N., Powis, K. & High, S. Post-translational translocation into the endoplasmic reticulum. Biochim. Biophys. Acta - Mol. Cell Res. 1833, 2403–2409 (2013).

35. Lakkaraju, A. K. K. et al. Efficient secretion of small proteins in mammalian cells relies on Sec62-dependent posttranslational translocation. Mol. Biol. Cell 23, 2712–2722 (2012).

36. Vilardi, F., Lorenz, H. & Dobberstein, B. WRB is the receptor for TRC40/Asna1-mediated insertion of tail-anchored proteins into the ER membrane. J. Cell Sci. 124, 1301–1307 (2011).

37. Stefanovic, S. & Hegde, R. S. Identification of a Targeting Factor for Posttranslational Membrane Protein Insertion into the ER. Cell 128, 1147–1159 (2007).

